# Responses to oddball communication sequences in the bat frontal and auditory cortices

**DOI:** 10.1101/2025.02.05.636670

**Authors:** Eugenia González-Palomares, Johannes Wetekam, Manuel S. Malmierca, Julio C. Hechavarria

## Abstract

Stimulus-specific adaptation (SSA) is a ubiquitous phenomenon in the animal kingdom across sensory modalities, but this type of neural response has rarely been studied using natural sounds in the auditory brain. Here, we leveraged the well-documented acoustic repertoire of the bat species *Carollia perspicillata* to study adaptation in the bat brain using natural communication sounds. We searched for SSA in single neuron spiking activity measured in two brain areas simultaneously: the auditory cortex and the frontal auditory field. The stimuli consisted of natural distress syllables, a form of vocalization produced by bats under duress. Bat distress vocalizations signal different degrees of urgency based on their amplitude modulation pattern, without large differences in their spectral structure. These distress vocalizations make an ideal test case for exploring the limits of neural deviance detection when considering naturalistic soundscapes with low stimulus contrast. The results show limited evidence for stimulus-specific adaptation in response to natural sound sequences in the majority of neurons studied. Many neurons did show a prominent effect related to context-dependent changes, caused by the type of sounds that occurred most frequently within each oddball sequence. Context-dependent responses were strongest in frontal neurons. Decoding analysis showed the existence of neural populations in both frontal and auditory cortices, which could distinguish between deviants and standards occurring within the same sequence, without large changes in evoked spike counts. Taken together, our results highlight the diversity of neural mechanisms complementing classical stimulus-specific adaptation when encoding natural vocalizations that do not differ markedly in their spectral composition.

## Introduction

The detection of unpredicted (or salient) stimuli is important for extracting changes in an animal’s environment. Efficient information coding in the nervous system devotes less resources to repeating than to important novel stimuli (Dong and Vicario, 2018; Font-Alaminos et al., 2021; Näätänen et al., 2007; Parras et al., 2017; Taaseh et al., 2011; Ulanovsky et al., 2003). A common way to study this phenomenon in the auditory domain is through the oddball paradigm, which consists of presenting a sequence of two sounds with different probability of occurrence: one of them presented often (standard) and the other one rarely (deviant). Studies using the oddball paradigm have demonstrated enhanced responses to rare deviants which occur alongside the suppressed responses to repetitive stimuli. These responses are defined as mismatch negativity if neural signals are recorded extracranially, or as stimulus-specific adaptation (SSA) if recorded from single neurons (Parras et al., 2017; Ulanovsky et al., 2003). SSA has been studied intracranially in multiple brain areas, including the inferior colliculus, auditory thalamus, auditory cortical fields and frontal areas, mostly using simple stimuli, such as pure tones (Antunes et al., 2010; Carbajal et al., 2024; Casado-Román et al., 2020; Duque et al., 2018; Gong et al., 2024; Hockley and Malmierca, 2024; Lao-Rodríguez et al., 2024, 2023; Malmierca et al., 2009; Parras et al., 2017; Pérez-González et al., 2024, 2021; Quintela-Vega et al., 2023; Ulanovsky et al., 2003; Wong et al., 2024; Yaron et al., 2020).

Recently, more complex sounds, including natural ones, have started to be used in studies of SSA and deviance detection in general (Harpaz et al., 2021; López-Jury et al., 2023; Wetekam et al., 2022; Yaron et al., 2020; Yoshino-Hashizawa et al., 2023). It has been rightfully argued that incorporating natural sounds is essential for understanding the biological value of deviance detection and its underlying mechanisms. Natural sounds occurring consecutively often overlap in their spectrum posing a challenge to classic SSA models that were developed for spectrally non-overlapping auditory stimuli (Hershenhoren et al., 2014; Mill et al., 2011; Taaseh et al., 2011). Our goal in this article is to assess whether SSA occurs in response to sequences of spectrally rich and overlapping communication vocalizations in a hearing specialist: the bat species *Carollia perspicillata*.

Both the auditory system and vocalizations of this bat species have been extensively described. *C. perspicillata* uses a broad repertoire of natural vocalizations to navigate and communicate socially (González-Palomares et al., 2021; Knörnschild et al., 2014; Thies et al., 1998). Among communication vocalizations, distress calls hold a unique and significant role. Distress calls are emitted under duress, for example, when a bat is captured by a predator (Klump and Shalter, 1984). These calls are commonly emitted in sequences composed of individual units (syllables) that have a broad spectrum (i.e., 20 to > 70 kHz) and high similarity between them (González-Palomares et al., 2021; Hechavarría et al., 2020). Despite the high similarity and repetitiveness observed within one distress sequence, transitions occur between fast amplitude modulated (AM) and slow-modulated syllables (Fig. 1). A single bat can produce the same syllable in both its slow and fast amplitude modulated forms (González-Palomares et al., 2021; Hechavarría et al., 2020). Amplitude modulations are a defining feature of agonistic vocalizations and they can be found in bat distress calls (González-Palomares et al., 2021; Hechavarría et al., 2020), rat 22-kHz vocalizations emitted during fear conditioning (Gonzalez-Palomares et al., 2023), and even in human screams (Arnal et al., 2015). In bats, listening to sequences of fast-AM distress calls increases the heart rate more than listening to sequences of slow-modulated vocalizations (Hechavarría et al., 2020). This finding indicates that amplitude modulations may signal higher levels of stress, a hypothesis supported by studies in rats subject to fear conditioning, in which the rats vocalized with more marked AMs during escape behaviors and in response to conditioned stimuli (Gonzalez-Palomares et al., 2023). In other words, detecting fast-AM vocalizations within streams of slow-modulated sounds could be of relevance to assess changes in the levels of stress by the emitter. In the wild, bats have to cope with transitions between slow and fast-AM vocalizations that do not differ markedly in their spectra (Fig. 1A). Yet, whether bat auditory neurons can detect such naturally occurring AM transitions remains unknown.

**Figure 1.**
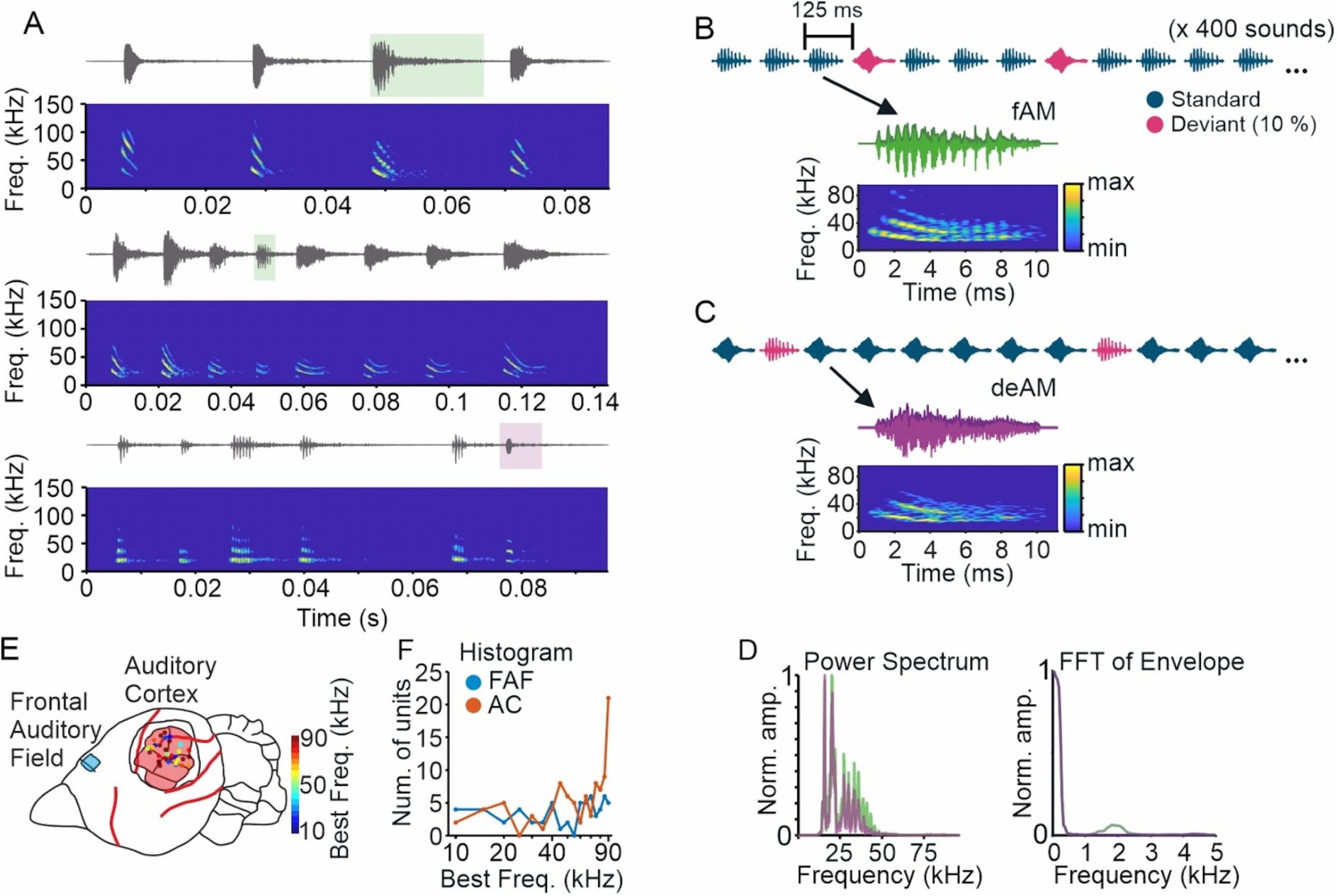
Examples of natural distress sequences and the oddball stimulation protocol used to study SSA. A) Oscillograms and spectrograms of three vocalization bouts recorded from conspecifics showing that they consist of both fast and slow AM calls. The position of deviant syllables is indicated with shaded squares in green if the deviant was fast modulated or in pink for slow modulated deviants. B-C) Schematic of the stimulation paradigm. Two stimulation sequences were used in which the roles of standard and deviant were swapped (B: deAM as deviant, C: fAM as deviant). Each of the sequence consists of 400 sounds composed of 360 standards + 40 deviants. Shown with the arrows are the oscillograms and spectrograms of the sounds used as deviant in each sequence. D) Power spectrum of the two vocalizations in B-C and their normalized FFT (fast Fourier transform) of the calls’ envelopes (secant method, temporal resolution: 0.05 ms). Note the bump of the fast AM call at ∼1.7 kHz. Spectrograms: 512 points, hamming window, 0.8 ms frame width and 0.05 ms frame shift. E) Schematic of the bat’s brain with the recorded areas in color (Frontal Auditory Field, FAF, in blue; Auditory Cortex, in red). The positions of the recordings in the auditory cortex are color-coded for the best frequency. F) Histogram of all the best frequencies of the units used in each brain region.

To address this question, we investigated SSA to sequences of slow and fast modulated distress syllables using parallel neural recordings with silicon probes from both the frontal and auditory cortices. Our results revealed a diverse pattern of neural adaptation, including classic, but also non-classic SSA, indicating that some neurons responded to standard stimuli with higher responses than to the novel deviant sounds. Upon further exploration of these complex responses, we found that the most repeated sound within an oddball sequence generally determines the level of neuronal responsivity. This, in turn, affects responses to both standard and deviant stimuli ultimately creating different types of neural categories according to their SSA patterns. Our results also uncovered that units whose firing increases with the repeated presentation of fast-AM syllables are more prevalent in frontal brain areas than in the auditory cortex (AC). Taken together, our results highlight the diversity of mechanisms that support the processing of naturalistic vocalization sequences in the mammalian auditory system.

## Results

When under duress, bats emit a wide variety of distress calls (González-Palomares et al., 2021; Hechavarría et al., 2020, 2016). Recordings from our bat colony demonstrate that these calls often include fast amplitude-modulated calls occurring alongside calls with less pronounced amplitude modulation within the same sequence (Fig. 1A, top and middle). Conversely, it is also observed that a call with slow AM can be emitted amidst more strongly amplitude modulated distress calls within the same bout (Fig. 1A, bottom).

Inspired by these behavioral observations, we set up an experiment aimed to understand how the brain of an awake listening bat processes transitions between fast-modulated and slow modulated distress vocalizations. Our underlying hypothesis was that the occurrence of deviant sounds (i.e. a fast AM vocalization surrounded by slow modulated vocalizations) would elicit a strong novelty response in the form of classic SSA. To test this hypothesis, we generated oddball sequences using a combination of two sounds as acoustic stimuli: 1) a fast-AM natural distress syllable (referred to here as fast amplitude modulated: fAM, Fig. 1B) recorded from a conspecific and 2) the same call but demodulated modifying its modulation power spectrum (deAM; Fig. 1C, see methods). These two vocalizations had similar spectra when considering their carrier frequencies but differed in their temporal modulation patterns (Fig. 1D).fAM and deAM vocalizations were then used to create oddball sequences consisting of 400 sounds, in which “standards” comprised 90% of the sounds, while the remaining 10% were deviants occurring in a pseudorandom order.

The experiments were repeated for another natural distress vocalization, as well as with AM and non-modulated 23-kHz pure tones, which correspond to the peak frequency of communication calls in this bat species (see Figs. S1 & S2, respectively). For each sound considered, there were altogether two sequences, one in which the fAM was the standard and deAM was the deviant, and a corresponding sequence in which the roles of deviants and standards were reversed. This approach allowed us to compare neural responses to the same sound when used as deviants vs. when it occurred as standard in the two different stimulation protocols. The sounds were presented at a rate of 8 stimuli per second. The recordings were conducted in fully awake animals with silicon probes inserted in the AC and frontal auditory field (FAF). Each probe recorded neural activity at depths of 0, 200, 800 and 1000 µm from the cortical surface.

Before acoustic stimulation with the oddball sequences, pure tones (ranging from 10 to 90 kHz in 5 kHz increments at 70 dB SPL) were played to characterize the tuning properties of the recorded units. Figure 1E shows the location of the two recorded areas in the bat brain with the recordings within the AC color-coded according to their best frequency. Most of the units in the AC had high best frequencies, typical from the dorsal AC fields that specialize in temporal computations and that respond strongly to communication sounds (Esser and Eiermann, 1999; Hechavarría et al., 2013; Martin et al., 2017). By contrast, the best frequencies were more evenly distributed in the FAF (Fig. 1F). Properties of the frequency tuning curves were further analyzed (Fig. S3) to check for correlations with other variables used in the upcoming sections of this manuscript (see below).

### Classic and non-classic SSA occur in auditory and frontal cortices

We recorded a total of 94 spike-sorted single units in the AC, and 60 units in the FAF. To compare differences in response to the same sound when it appeared as standard or as deviant, the stimulus-specific adaptation index (SI) was computed (Ulanovsky et al., 2003). This index is calculated as:

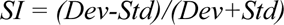

where *Dev* and *Std* are the number of spikes fired by the units in response to the same sound when it occurred as deviant or standard, respectively. Note that positive SIs indicate stronger responses to the same sound when it was presented as deviant versus when it was presented as standard (that is, classic deviant detection), while negative values indicate the opposite pattern. Figure 2A-B illustrates the distributions of SIs calculated for AC and frontal areas. The inspection of the distributions allows to readily observe that SIs values cluster towards the center of the graph, with no particular preference for any of the quadrants that indicate different types of SSA in both brain areas studied (centroid of AC: 0.001, -0.003; FAF: -0.011, 0.013). For example, in the AC, the number of units that displayed classic SSA (SI>0) in response to both the fAM and deAM sounds (quadrant I, n=19 units) was qualitatively similar to the number of units with non-classic SSA (SI<0) to both stimuli (quadrant III, n=21 units). Classic SSA has been typically found in studies using pure tones that do not overlap in their spectra, and it indicates consistently stronger responses to a sound when it is used as “deviant” vs. when the same sound is the “standard” under the oddball paradigm stimulation. In seminal SSA studies, SI distributions are typically shifted towards quadrant I, but our results show that this is not the case when using spectrally overlapping vocalizations that differ only in their AM pattern. These results were apparent in the AC (Fig. 2A), as well as in the FAF (Fig. 2B), although in the latter, more neurons were located in quadrants II and IV, indicating classic SSA to one stimulus but not to the other one.

**Figure 2.**
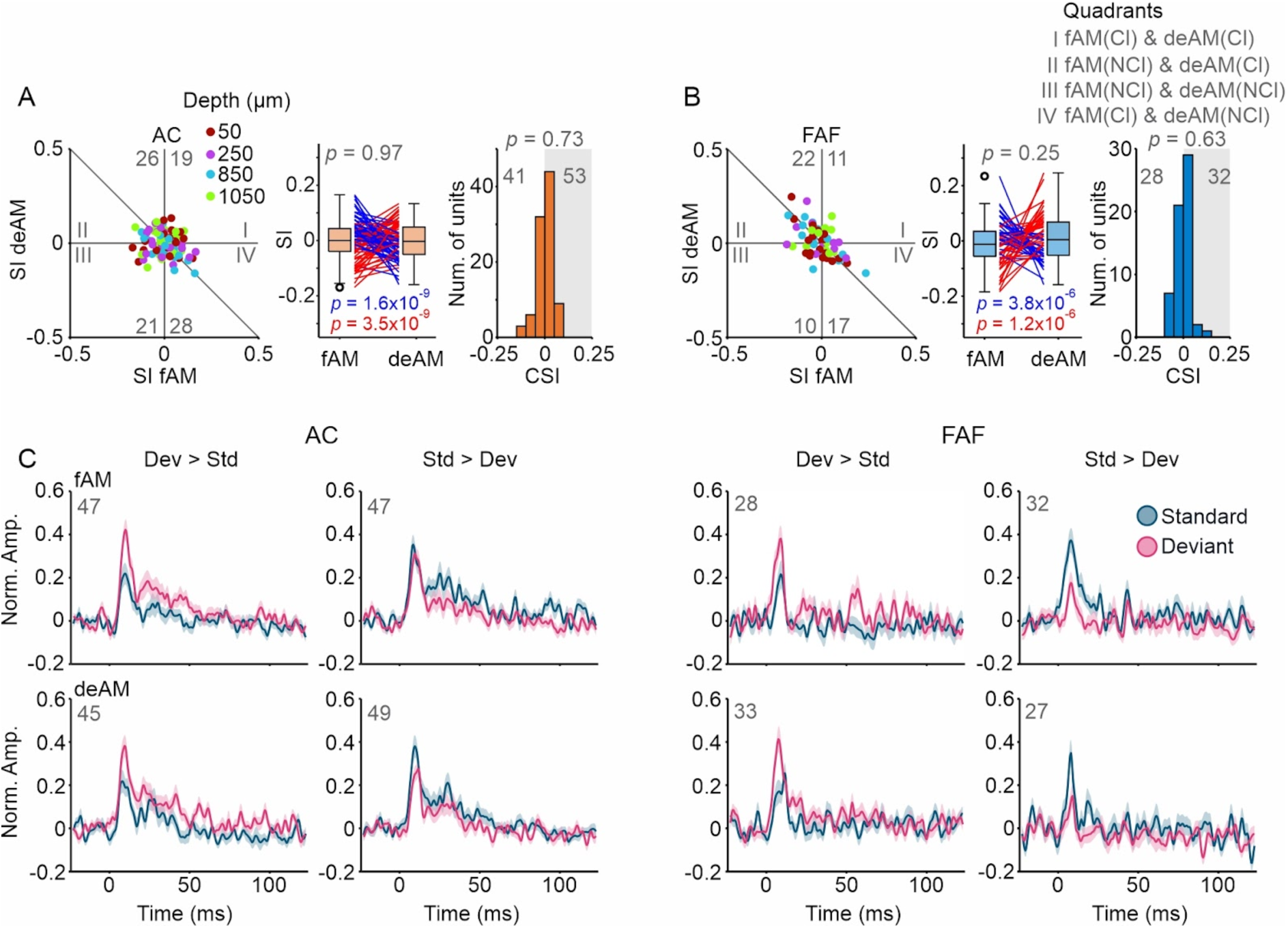
Neural activity in response to the oddball sequence. A) Scatter plot (left) of SI for both stimuli (fAM, x axis, deAM, y axis) color-coded for the depth for recordings in the Auditory Cortex (AC). The number of the units in each quadrant is indicated in gray. Box plot (middle) with lines of all the SI of fAM and deAM of all units. Blue lines: SI fAM > SI deAM, and red lines, SI fAM < SI deAM. *P* values of the Wilcoxon signed-rank test of the SI of both stimuli (in gray for all units, blue for units with “SI fAM” > “SI deAM”, red for “SI fAM” < “SI deAM”). Histogram (left) of common-SI (CSI). The number of the units on each side of the histogram is given in gray, and the shaded area represents CSI > 0. Cl: Classic (SI > 0), NCl: Non-classic SSA (SI < 0). B) Same as A, for Frontal Auditory Field (FAF). C) Mean ± SEM (standard error of the mean) of probability density functions (Gaussian kernel of width 1 ms, at steps of 0.5 ms) during the deviant (pink) responses to the sounds (fAM, top row; deAM, bottom row) and the standard (dark blue, using the same number of sound presentations as for the deviant N = 40 trials).

When we compared the SI indices obtained with the two sounds used (fAM and deAM) we observed no significant differences in either brain area (Wilcoxon signed rank test, *p*= 0.97 for the AC and *p* = 0.25 for the FAF, see Fig. 2A & B middle panels). However, we did notice an interesting trend in which units that had a larger SI in response to the fAM stimulus exhibited lower SI values for the deAM sound, and vice versa. The latter occurred in both brain areas (blue and red lines in Fig. 2A & B middle panels). This emphasizes the differing SSA behavior of AC and frontal neurons when listening to oddball sequences that alternate between amplitude modulated and demodulated vocalizations despite the lack of large spectral differences. We also examined the distributions of the common-SI (CSI, see Methods). In both areas there were slightly more units with positive CSI, which indicate overall stronger responses to the deviants. However, at the population level, the CSI distributions’ medians were not significantly different from zero (AC: median= 0.006, *p*=0.73; FAF: median=0.003, *p*=0.63, Wilcoxon signed rank test).

Overall, the results described above indicate the existence of at least two types of neurons: 1) those that show stronger responses to deviant sounds (classic SSA) and 2) those that showed stronger responses to repetitive standards (non-classic SSA behavior). To confirm the existence of these two neural types, average population responses were plotted separately for each group of neurons in the two brain areas studied, and for each of the two stimuli used (Fig. 2C, see also Fig. S1 and S2 for similar results obtained with different stimuli). The average response plots show that both brain areas studied (AC and FAF) displayed strong responses to the sounds used as stimuli. As expected, classic SSA involved stronger responses to the deviant than to the standard sound, regardless of the stimulus to which they occurred (classic SSA: pink line is higher than the dark blue line in Fig. 3C). On the other hand, non-classic responses were related to higher responses to the standard than to the deviant sound (non-classic SSA: dark blue line is higher than pink line in Fig. 3C). These observations confirm the existence of both classic and non-classic SSA in both the AC and FAF of awake bats when listening to sequences of amplitude modulated and demodulated vocalizations that do not differ largely in their power spectrum.

**Figure 3.**
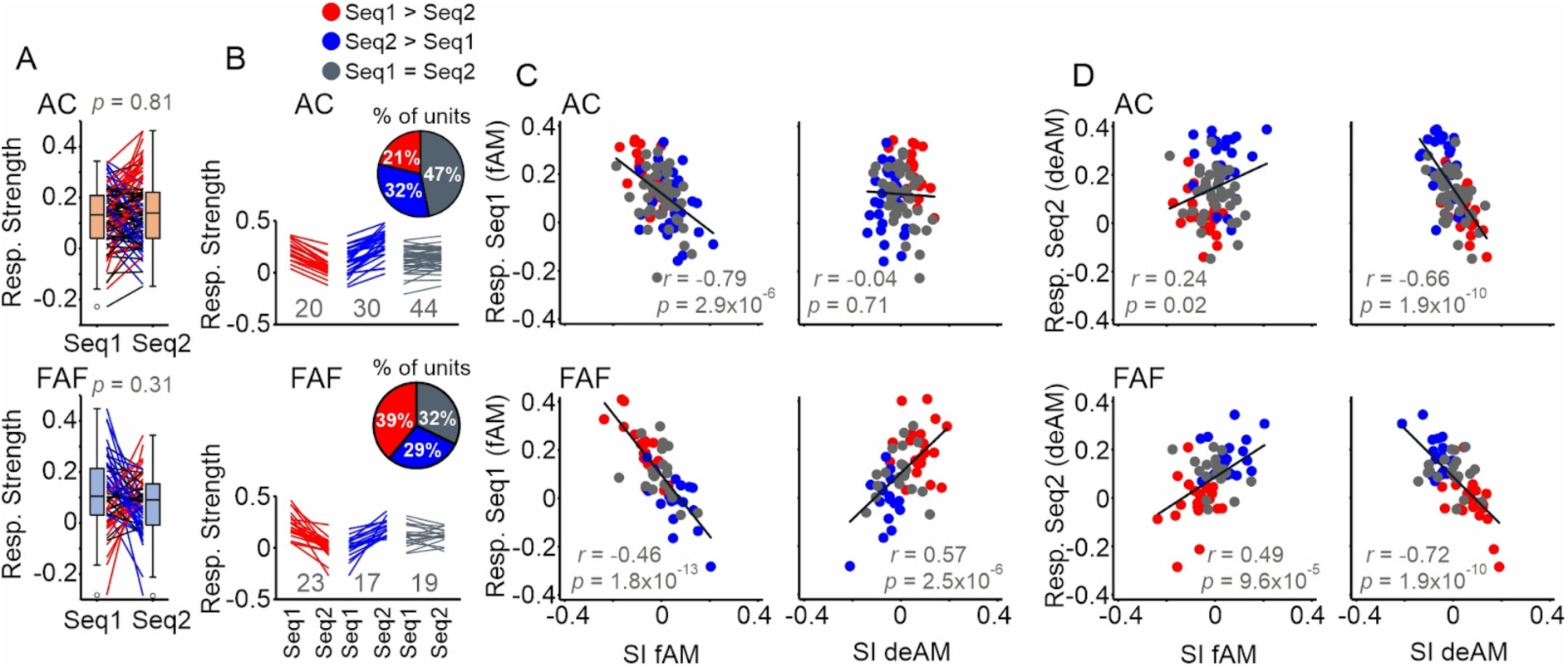
Exploring the relation between response strength and SI. A) Response strength (mean of activity response of forty standards) for each sequence (Seq1: fAM as Standard; Seq2: deAM as Standard). There were no significant differences between the response in the two sequences (Wilcoxon signed-rank test) when all the units were considered. We further classified units classified into three groups, in red (significant response difference between Seq1 and Seq2, Wilcoxon rank-sum test, *p*<0.05, with response in Seq1 > Seq2), in blue (same as for the red group, but with response in Seq2 > Seq1), and gray (no significant difference between Seq1 and Seq2). Each line represents a unit, with the total number of units shown below. The plots are shown for the AC (top) and FAF (bottom). Insets show the percentage of units in each group. B) Line plots showing the same as in A, but for all groups separated. C) Scatter plots of the response strength in Seq1 and the SI of both stimuli fAM (left) and deAM (right) for AC (top) and FAF (bottom). Linear regression fit (gray line) for all units, with the Pearson’s correlation coefficient and its corresponding *p* value also shown. D) Same as in C for Seq2.

In our data, SI was not positively correlated with the properties of the frequency tuning curves (Fig. S3), nor did it depend on the spike shapes of the recorded neurons (Fig. S4). We also evaluated the spontaneous activity prior to the acoustic stimulations and found no significant correlation with the SI (Fig. S5). In addition, to investigate the similarity in responsiveness between simultaneously recorded units, we analyzed all possible pairs of neurons both within and between brain areas (Fig. S6). We found that the highest probability of finding matching pairs (both units being either Classic or Non-classic SSA) occurred in the FAF for both stimuli, followed by the simultaneously recorded units in FAF and AC for the fAM stimulus. In the other conditions, mismatched pairs comprised about half of the pairs and in most cases the probability of finding matching pairs of neurons did not differ from a randomly generated control distribution (see Fig. S6C).

### Repetitive acoustic context alters neural responsivity

Previous research that reported inconsistent classification of units based on their SSA responses suggested that these neural patterns might arise from changes in the spike rate throughout an entire oddball sequence without necessarily being linked to the occurrence of deviant sounds (Harpaz et al., 2021). In our data, overall activity levels could vary as certain natural sounds may influence neural responsivity more than others (i.e., a neuron could preferentially respond to fAMs or deAMs). Therefore it could be misleading to compare responses obtained when the same sound is used as standard and deviant, as these responses arise from different stimulation sequences presented at different times potentially with varying underlying activity levels driven by the most repeated sound within each sequence. A similar scenario was found to occur in previous studies when using the so-called “tone-clouds” (Harpaz et al., 2021). Tone-clouds are spectrally complex and overlapping stimuli (like the vocalizations used here) but they lack ethological meaning for the animals.

We tested whether a similar phenomenon occurred in our data obtained with fast AM and demodulated vocalizations. To do this, we looked into overall levels of responsivity by averaging the responses to 40 standard presentations of the same sequence (named Seq1: fAM as Standard and deAM as Deviant, and vice versa for Seq2). When considering all units, there were no statistical differences in the overall response strengths between the sequences (Fig. 3A, Wilcoxon signed rank test). Upon examination of individual units more closely, we identified three groups of neurons: those that responded stronger to Seq1 (red in Fig. 3A; Wilcoxon rank sum test, alpha = 0.05), those that responded stronger to Seq2 (blue), and those that showed no statistical difference between the two sequences (gray). About half of the units in the AC showed no differences in response to either sequence, while the remaining units were equally split, with some units responding more to Seq1 and others to Seq2 (Fig. 3B, top). In the FAF, the largest group of units was more responsive during Seq1. It is important to note that Seq1 contained more fast amplitude modulated vocalizations (fAM is the standard sound in this case). Further, as mentioned in the introduction, fAMs are typically used to signal higher levels of urgency in bats and other species (Arnal et al., 2015; Gonzalez-Palomares et al., 2023; González-Palomares et al., 2021; Hechavarría et al., 2020). We also checked for correlations between response strength and spontaneous activity and the only statistically significant correlation occurred during Seq2 (deAM as Standard) in the AC (Fig. S5B). Simultaneously recorded pairs of units in the AC had significant correlation in the responsivity but only during Seq1 (Fig. S6D).

To better evaluate the relationship between sequence-evoked responsiveness and SI, we correlated both parameters across all neurons for both stimuli used (Fig. 3C for Seq1; Fig. 3D for Seq2). We found that, in almost all cases, responsiveness was correlated with SI across all neurons, with some correlations being negative and others positive. Additionally, we observed that correlations obtained with different stimuli followed opposite patterns. The data show that neurons with a preference for a given stimulus – quantified by stronger responses to the sequence where this stimulus was the standard – tended to have a more negative SI for this preferred stimulus (indicating non-classic SSA behavior) and a more positive SI for the unpreferred stimulus (indicating classic SSA; Fig. 3C and D, right panels). This can occur because the repetitive stimulation with the preferred stimulus increases the overall response to all the sounds in that sequence (including the rarely happening deviants), leading to more negative SI values for the preferred, and more positive values for the unpreferred stimuli of a given neuron. This suggests that general changes in responsivity to the stimulation sequences can lead to SSA-like effects in neurons without necessarily involving deviance detection.

### Different response patterns enable discrimination of co-occurring modulated and demodulated vocalizations

The results described above indicate that changes in overall responsivity levels may resemble SSA effects. This makes it challenging to compare responses across stimulation sequences in the way conventionally done in SSA studies using artificial stimuli. However, this does not necessarily mean that individual neurons cannot display different responses to fAM and deAM sounds that co-occur within the same oddball sequence. To test this idea, we trained support machine (SVM) classifiers with single trial spike patterns obtained in response to standard and deviant sounds presented within the same oddball sequence. The goal was to quantify how many neurons could distinguish between responses to fAMs and deAMs presented within the same sequence. It is important to note that successful discrimination using classifiers does not necessarily rely on an increase in spike count, but could also occur based on a differential distribution of spikes over time.

The classifiers were trained with single-neuron data (40 training trials: 20 standards, 20 deviants). Another 40 trials were selected as testing dataset. This procedure was repeated 50 times for each neuron, with the training and corresponding testing datasets randomly selected in each iteration. The obtained classification accuracy was compared with data obtained by shuffling the class identities in the training dataset (Fig. 4; see also Figs. S1G and S2G for similar analysis with data obtained in response to two other pairs of stimuli). In the AC, we observed that 37 and 41% of the neurons showed significant classification performance for sequences 1 and 2, respectively (35/94 and 39/94 for Seq1 & Seq2). In other words, these are the neurons whose response could be used to discriminate between fAM and deAM sounds that co-occurred in the same oddball sequence. In the frontal cortex, the percentage of neurons able to discriminate between co-occurring fAMs and deAMs was higher than in the AC, reaching 53% (31/59 neurons) and 59% (35/59) for Seq1 and Seq2, respectively.

**Figure 4.**
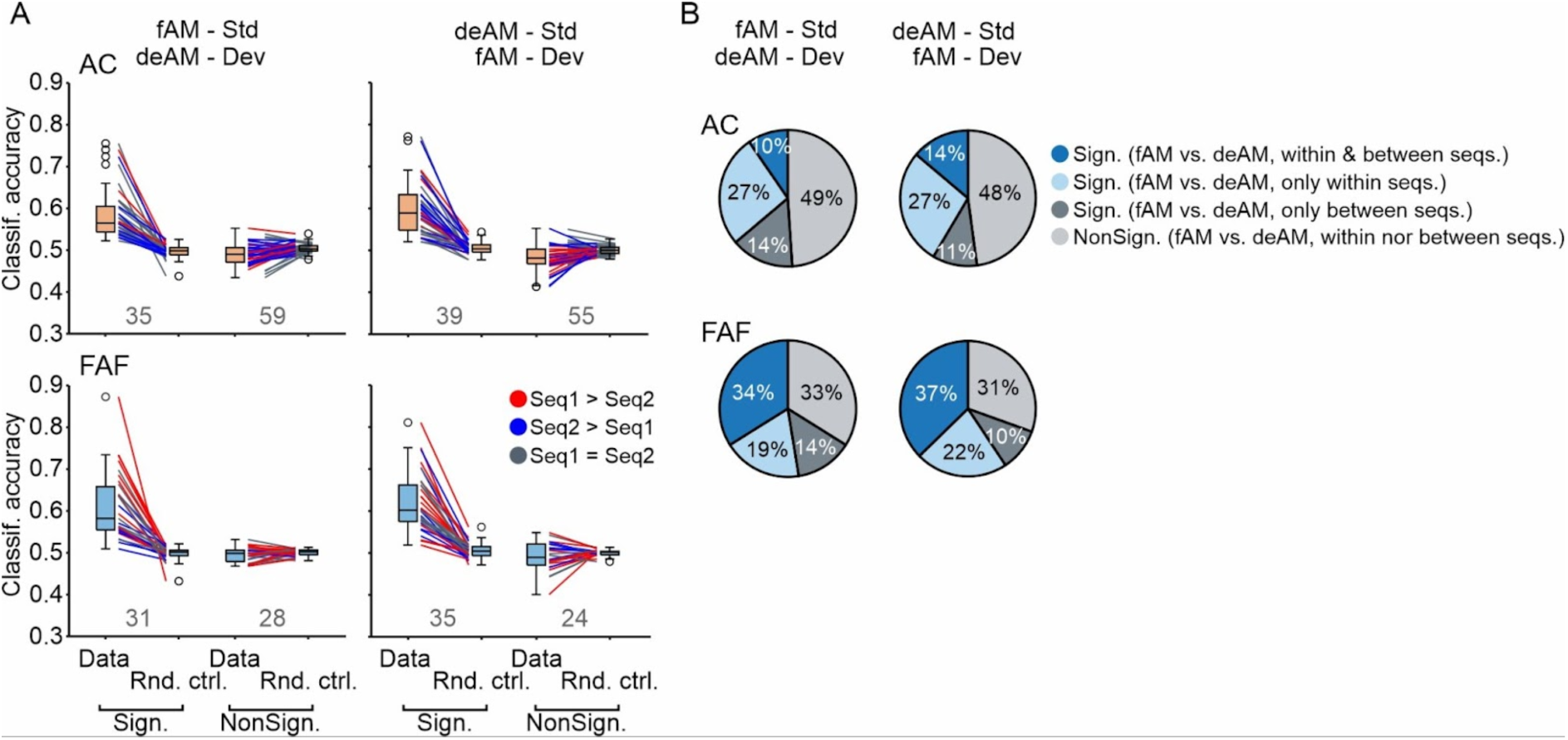
Box plots of the prediction accuracy calculated using binary SVM classifiers trained and tested with randomly chosen single trials (20 standards and 20 deviants for training + another 20 standards and 20 deviants for testing). The analysis and statistics was done for each unit by generating 50 SVM models each time with randomly chosen test training and testing datasets. For the random control, SVM models were generated again in a similar manner but with shuffled “standard”/”deviant” labels. Shown in the figure are the mean of the prediction accuracies for each unit. The units have been separated by whether they could significantly tell apart standards and deviants that co-occurred within the same sequence after comparing the results of the 50 iterations with the shuffled control (Wilcoxon signed-rank test, *p*<0.01). This was done for the AC (top) and FAF (bottom) and for stimulation sequences 1 (left) and 2 (right). The number of units within each group (i.e. neurons that predict and neurons that do not) is given with grey numbers. Lines are colored using the same convention as in Fig. 3, to indicate whether they had a preference for the sounds that were more repeatable in one sequence or another. B) Pie charts indicating the results obtained when using SVM models trained with data from standard sounds from both sequences. This analysis was done to evaluate if the units could also be able to discriminate between fAM and deAM sounds in different contexts. Dark blue slices shows the proportion of units that have significant prediction accuracies in both cases (fAM vs deAM within the same sequence, and fAM and deAM in different sequences), the light blue slice units that are significant within the same sequence but not between sequences, in dark gray, units that were only significant between the sequences, and in light gray, units that were non-significant prediction accuracies within the same sequence.

In summary, our results indicate that context effect interferes with SSA estimation when considering responses across oddball sequences (see previous section). However, analyzing responses within individual stimulation sequences does show the existence of neural populations whose firing differentially represents the spectrally overlapping fAM and deAM vocalizations. This neural population is not negligible, as it comprises >37% of the neurons in both the AC and frontal cortex. Out of the neurons that could discriminate between co-occurring fAM and deAM vocalizations, 93% (130/140, across all combinations) showed no significant differences in spike count between the deviant and standard trials of the same sequence (using a bootstrapping test, Fig. S7). This suggests that the discrimination ability does not rely on a change in response strength, but rather on the timing of evoked spikes. When comparing the response to the same sound presented as standard or as deviant (i.e., in different sequences), 58% (31/59 and 37/59 for fAM and deAM, respectively) of the FAF neurons had significantly different responses, and 30% (20/94 and 37/94 for fAM and deAM, respectively) of the AC neurons.

A logical question to ask is whether the neuronal population that could distinguish co-occurring fAMs and deAMs in an oddball sequence could also do so if the two sounds occurred outside of the oddball context. To test this idea, we trained a new set of SVM classifiers, this time considering only responses obtained when the fAMs and deAMs were presented as standards in the two oddball sequences. The goal was to identify neurons that differentiate between sounds when they are repeated multiple times without any novelty. In both the AC and FAF, we found a subset of neurons that could consistently distinguish between fAMs and deAMs both between and within sequences (dark blue in the pie charts in Fig. 4). This population was most prominent in the frontal part of the brain indicating that frontal neurons exhibit a stronger baseline fAM-vs.-deAM selectivity when compared to AC neurons. Additionally, 27 % of AC neurons could discriminate fAM-vs.-deAM only within the oddball sequences, suggesting that these neurons are sensitive to stimulus probability differences (deviance detection, but not based on an increase in spike count). This type of neuron was more prominent in the AC than in the frontal cortex (light blue in the pie charts in Fig. 4). Between 10 and 14 % of neurons could distinguish the vocalizations types only in different sequences (dark gray). The response of the remaining neurons (light gray in the pie charts in Fig. 4) could not differentiate fAMs from deAMs, neither when they co-occurred as standards in different sequences, nor when they co-occurred as standard and deviants in the same oddball sequence. Taken together, our results demonstrate that about half of the neurons studied in the AC and FAF can detect transitions between deAM and fAM vocalizations but only between 19 and 27% of the neurons gain novelty detection within oddball sequences.

## Discussion

In this study, we recorded from the auditory cortex and frontal auditory field of awake bats (*C. perspicillata*) while they listened to oddball sequences comprising natural amplitude-modulated (AM) vocalizations (recorded from a conspecific) and their demodulated versions. Our findings reveal global changes in neural responsivity, which appear to be linked to the most repeated stimulus in the oddball paradigm (i.e. the standard sound). Such context-dependent responses can sometimes masquerade as classic SSA and non-classic SSA responses. Despite these context-dependent changes in neuronal responsivity, we show that over 40% of the neurons in both the AC and FAF could distinguish between modulated and demodulated vocalizations that co-occurred within the same oddball sequence. Nearly 20% of the neurons demonstrate this discrimination ability only when amplitude modulated and demodulated vocalizations co-occurred within the same oddball sequence, but not between sequences, indicating that these neurons encode stimulus probability. Our results highlight the challenges of studying stimulus-specific adaptation using natural vocalizations with ethological relevance but also overlapping spectra. While studying responses to conspecific vocalizations brings us one step closer to understanding how the brain operates in natural conditions, it also requires the use of more sophisticated tools that go beyond simple spike counts for quantifying responses.

Previous SSA studies in bats have shown differences in the processing of novelty detection for natural echolocation and communication calls in the auditory cortex (López-Jury et al., 2023, 2021) and in subthalamic stations (Wetekam et al., 2024). Deviance detection in bats tends to favor echolocation deviants when presented within a communication context. This highlights the importance of the ethological relevance of the sounds used for stimulation, which is crucial for our understanding of how the brain processes novel stimuli (Wetekam et al., 2024). Although previous work has provided indisputable evidence of SSA in bats, it is important to consider that echolocation and communication sounds differ markedly in their spectra, in the same way pure tones used in classic SSA studies do. Amplitude-modulated distress calls paired with their unmodulated versions are part of the natural bat soundscape, with bats alternating between these two sound types in natural distress sequences (see Fig. 1M; González-Palomares et al., 2021; Hechavarría et al., 2020). Studying SSA with both modulated and demodulated vocalizations challenges existing SSA models. While modulated and demodulated vocalizations are very similar in their spectra, the presence of a modulating wave increases the peakedness (or harmonic-to-noise ratio) of AM sounds (Hechavarría et al., 2020).

The main synaptic model used to explain SSA evoked by spectrally non-overlapping stimuli is based on convergent inputs tuned to different frequency bands (Hershenhoren et al., 2014; Mill et al., 2011; Taaseh et al., 2011). These inputs, which differ in their frequency tuning profile, enable adaptation to one sound (i.e., the standard) while the other input remains unadapted and ready to respond when a deviant occurs. Such a model struggles to produce SSA in response to sounds with overlapping spectra, such as the natural vocalizations used in this study, or the artificial tone clouds used in previous work (Harpaz et al., 2021). When sounds overlap in their spectra, both deviants and standards would activate the same inputs and SSA cannot occur unless the convergent inputs are selectively “tuned” to the fAM and deAM vocalizations. Our data suggest that some neurons in the AC and FAF preferentially respond to either modulated or demodulated vocalizations, regardless of context (∼12% in the AC and ∼35% in the FAF, see Fig. 4). However, the majority of cells responded equally well to both sound types. This indicates that there are no major, selectively tuned pathways that respond exclusively to one or the other type of vocalizations.

In approximately 20% of cells we observed an increase in discriminability between fAM and deAM vocalizations presented within the same oddball sequence. It is important to note that an increase in discriminability does not necessarily imply stronger responses to the deviant sound (as stipulated in classic SSA theory), but it does indicate that some individual neurons are capable of discriminating transitions between fAMs and deAMs sounds within the same acoustic stream. In both the AC and FAF, approximately 40% of the neurons detect such transitions, when considering both those that gained discriminability in the SSA context and those that retained their discriminability from across-sequence comparisons. Note that we assessed differences between fAM- and deAM-evoked responses by using SVM classifiers, which can detect subtle differences in the temporal pattern of single-trial spiking responses. Such small differences could have been overlooked if only overall spike counts were considered. Differences in spike patterns to co-occurring deviants and standards do not necessarily require segregated inputs that converge into the same neuron. They rather could be based on activity driven by the same inputs and the subtle deviations on excitability created by the preceding sounds.

In the frontal cortex, we observed that repetitive sequences could either increase or decrease the spikiness of many cortical neurons. One possible explanation for these global changes in responsivity is related to the sidebands and peakedness created by the amplitude modulations. As mentioned above, the presence of a modulatory wave increases the peakedness of the spectrum and alters the harmonic to noise ratio of natural vocalizations (Walker et al., 2011). Such changes do not affect the overall spectra of the calls in a way that would cause modulated and demodulated vocalizations to activate different inputs to cortical neurons. However, changes in spectral peakedness could still lead to changes in responsivity to the sequences based on different levels of activation in the same inputs to the studied neurons. Another possibility could be tracking amplitude modulation cycles in the calls (Johnson et al., 2012; Joris et al., 2004), although this seems unlikely since bat vocalizations are modulated at 1.7 kHz (González-Palomares et al., 2021; Hechavarría et al., 2020), a rate too fast for locking spikes to individual AM cycles. Although auditory nerve fibers can phase lock up to 5 kHz in cats (Johnson, 1980), the ability to use the temporal coding mechanism decreases along the ascending auditory hierarchy (Joris et al., 2004). In fact, the majority of AC neurons of *C. perspicillata* cannot track amplitude modulations faster than 20 Hz (García-Rosales et al., 2018; Martin et al., 2017).

Taken together, the results presented in this manuscript provide an account of how deviance detection could operate in oddball sequences of spectrally complex and overlapping natural vocalizations. Some neurons appear to detect such low-contrast deviants, but they are challenging to identify, as changes in spikiness are rather subtle. Future studies including larger neuronal sample sizes and testing other sound types beyond distress calls will help determine whether these findings are generalizable. Our results showing strong context-dependent changes in neural responsivity, align with previous studies in rodents that used artificial tone-clouds as stimuli and reported differentially enhanced responsivity to sound sequences (Harpaz et al., 2021). We present evidence from two cortical areas of an auditory specialist, demonstrating that enhanced neural responsivity and within-sequence discrimination of nearly identical sounds can co-occur when listening to natural vocalization streams.

## Methods

### Animals

Five male adult bats (species *C. perspicillata*) were used with initial weights 21.4 ± 1.34 g (mean ± STD). The animals were taken from the bat colony at the Institute for Cell Biology and Neuroscience at the Goethe University in Frankfurt am Main, Germany. The colony is in a temperature- and humidity-controlled room with *ad libitum* access to food and a 12h light/dark cycle. The experiments comply with all current German laws on animal experimentation. All experimental protocols were approved by the Regierungspräsidium Darmstadt, permit #FU-1126. The study was conducted in accordance with the ARRIVE (Animal Research: Reporting of *in Vivo* Experiments) guidelines.

### Surgery and electrophysiological recordings

Bats were anesthetized subcutaneously with a mixture of ketamine (10 mg/kg Ketavet, Pharmacia GmbH, Germany) and xylazine (38 mg/kg Rompun, Bayer Vital GmbH, Germany). Local anesthesia (ropivacaine hydrochloride, 2 mg/ml, Fresenius Kabi, Germany) was injected subcutaneously on the skull before surgery and whenever the wounds were handled. The bats were placed in a custom-made holder that was kept at 28°C with a heating pad. The skin and muscle covering the skull were removed. Subsequently, a custom-made metal rod (1 cm long, 0.1 cm diameter) was glued onto the skull using acrylic glue (Heraeus Kulzer GmbH), super glue (UHU) and dental cement (Paladur, Heraeus Kulzer GmbH, Germany). Two craniotomies were performed in the left hemisphere using a scalpel blade around the regions of interest: the auditory cortex and frontal auditory cortex (see papers for more details), in each case the brain surface exposed was ∼ 1 mm^2^.

The bats were left at least two days for recovery before the recordings, which were performed on non-consecutive days, not for longer than 4 h and not later than 14 days after the surgery. The bats were euthanized with an anesthetic overdose (0.1 ml pentobarbital, 160 mg/ml, Narcoren, Boehringer Ingelheim Vetmedica GmbH, Germany). The animals received water every 1.5 h.

The recordings were done in an electrically shielded and sound-proofed Faraday cage. The bats were head-fixed in the custom-made holder. The recordings consisted of a maximum of two protocols: iso-level frequency tuning and oddball-paradigm with a natural and a demodulated sound. On the first recording day, two small incisions in the skull were made for the short-circuited reference and ground wires in non-auditory areas. The recording electrodes (A1×8-4mm-200, NeuroNexus, Ann Arbor, MI) had 8 channels with 200 µm inter-site spacing. Each electrode was introduced in a region of interest, perpendicular (AC) and tangential (FAF) to the brain surface until all the channels are in the tissue with help of Piezo manipulators (PM-101, Science products GmbH, Hofheim, Germany). The electrophysiological signals were amplified (USB-ME16-FAI-System, Multi Channel Systems MCS GmbH, Germany) and stored in a computer using a sampling frequency of 25 kHz. The data were stored and monitored on-line in MC-Rack (version 4.6.2, Multi Channel Systems MCS GmbH, Germany).

### Acoustic stimulation

We used three protocols here: pure tones for iso-level frequency tuning, oddball paradigm with natural calls and oddball paradigm with amplitude modulated pure tones. Sounds were played from a sound card (ADI-2-Pro, RME, Germany) at a sampling rate of 192 kHz, connected to a power amplifier (Rotel RA-12 Integrated Amplifier, Japan) and to a speaker (NeoX 1.0 True Ribbon Tweeter; Fountek Electronics, China) that was about 15 cm away from the right ear. The speaker was calibrated using a microphone (¼-inch Microphone Brüel & Kjær, model 4135) recorded at 16 bit and 384 kHz of sampling frequency with a microphone amplifier (Nexus 2690, Brüel & Kjær).

For the frequency tuning, the pure tones (10 ms duration, 0.5 ms rise/fall time) of frequencies from 10 to 90 kHz in steps of 5 kHz at a level of 70 dB SPL. The pure tones were played in a pseudo-random manner with a total of 20 repetitions per sound.

For the oddball paradigm with natural calls, we used natural calls recorded from a bat in our colony which are fast amplitude modulated (González-Palomares et al., 2021; Hechavarría et al., 2020). These calls were demodulated (Hechavarría et al., 2020). Each call was paired together with its demodulated version for the presentation in the sequences. Each sequence consists of 400 sounds in total, 90% of which correspond to one of the two calls (standard) and the remaining 10%, its pair, the order in the sequence is random. In the complementary sequence, these calls have their roles (standard and deviant) switched. Therefore, there were four sequences in total presented at random order. The calls within a sequence are presented at a rate of 8 Hz (every 125 ms) and there was an inter-sequence time of around 2 min.

### Spike detection and sorting

We used SpykingCircus (median absolute deviation of 5; templates of 3 ms) in order to detect and sort spikes (Yger et al., 2018). For each electrode, the cluster with more spikes was selected. For the frequency tuning curves, it was considered the spikes occurring during the 110 ms after stimulus onset.

### Analyses

Due to technical issues, channels at depths of 400 and 600 µm in both electrodes were not taken into consideration for further analyses. Units were considered to be responsive then they fire at least 1 spike in 10% of the trials of a sequence for both sequences of each stimulus pair, and additionally that is statistically significant (t-test, 0.05 significance level) the numbers of spikes fired in the first 25 ms post onset compared to 25 ms before the onset of the same presentations.

The peristimulus time histograms (PSTHs) shown in Figure 2 are probability density functions of the spike times considering 40 trials (Deviant or Standard), using a kernel smoothing function (Gaussian window of 1 ms bandwidth at 0.5-ms steps), that have been baseline-demeaned (mean value during the 23 ms before stimulus onset) and normalized to the maximum PSTH value of all four conditions: fAM as Standard, fAM as Deviant, deAM as Standard, deAM as Deviant). This was done for all units, and the mean ± SEM is what is shown in Fig. 2C. Following these, the stimulus-specific adaptation indices (SIs) for each stimulus were calculated as follows (Ulanovsky et al., 2003): (Dev - Std)/(Dev - Std), where “Dev” and “Std” correspond to the sum of the values in the first 25 ms after stimulus onset, for each condition respectively. Similarly, the common-SI (CSI; (Ulanovsky et al., 2003)) was calculated: (Dev_fAM_ + Dev_deAM_ - Std_fAM_ - Std_deAM_)/(Dev_fAM_ + Dev_deAM_ + Std_fAM_ + Std_deAM_). For the deviant sounds, the 40 presentations were considered, and for the standard, the 40 presentations that occurred before the deviants (or presentations randomly selected in the cases when that was not possible). For the spontaneous activity, it was considered the average spiking rate of 5 windows of 2-s duration randomly selected from the 15 s before the sequence onset.

To compare how alike the two recorded areas are while being recorded simultaneously, we evaluated the probability of finding a matching pair (pair of units that were Classic & Classic or Non-classic & Non-classic) both within and between areas (Fig. S6). This was done by taking 100 pairs from the real data (simultaneously recorded) and comparing them with randomly combined units (not necessarily simultaneously recorded). This process was repeated 1000 times for each condition and the probabilities of finding a matching pair was compared using the Wilcoxon rank sum test and the Cliff’s delta (Romano et al., 2006).

In order to assess if the units could discriminate between the standard and deviant sounds, we trained support vector machine (SVM) classifiers using single-trial spiking patterns presented within the same sequence. For the training set, 40 trials were selected (20 standards, 20 deviants), and another 40 different trials were used for the testing dataset. This process was repeated 50 times for each neuron, with the training and testing datasets being randomly selected each time. The classification accuracy results were compared to those obtained by shuffling the class identities (control condition) in the training dataset (Fig. 4). Units that had significantly higher prediction accuracy results (one-sided Wilcoxon signed-rank test, *p*<0.01) than using shuffled trials were shown separately. SVM models were also used in the same manner but comparing the responses to standards in one sequence to the responses to the standards in the other sequence to determine if the classifier could distinguish different sounds that have the same probability of appearance. For the training datasets, 20 standards from one sequence and 20 standards of the other sequence were used, and for testing, the same number of trials from different presentations. The process was repeated 50 times for each unit and with shuffled labels for the control condition.

### Statistics

All statistical tests were performed using custom-written Matlab scripts (R2023a, MathWorks, Natick, MA). When distributions were not normal, determined using the Kolmogorov-Smirnoff test, non-parametric tests were used, Wilcoxon rank sum, and the Wilcoxon signed rank, for paired comparisons. Otherwise, t-tests were used.

## Code availability

The scripts are available from the corresponding authors on reasonable request.

## Data availability statement

The original data of this study are available from the corresponding authors upon reasonable request.

## Acknowledgements

We thank our funding sources: Deutsche Forschungsgemeinschaft project # 275755787 (to J.C.H.); Spanish PID2023-148541OB-I00 funded by MCIN/AEI/10.13039/501100011033; Foundation Ramón Areces grant # CIVP20A6616 and Consejería de Educación, Junta de Castilla y León grant # SA218P23 (to M.S.M.).

## Author contributions

All authors designed the study; E.G.P. carried out the experiments; E.G.P. analysed the data; E.G.P. wrote the first draft of the manuscript; all authors revised the manuscript.

## Competing interests

The authors declare no competing interests.

## Notes

### Competing Interest Statement

The authors have declared no competing interest.

