## Supplementary Figures for "Responses to oddball communication sequences in the bat frontal and auditory cortices"

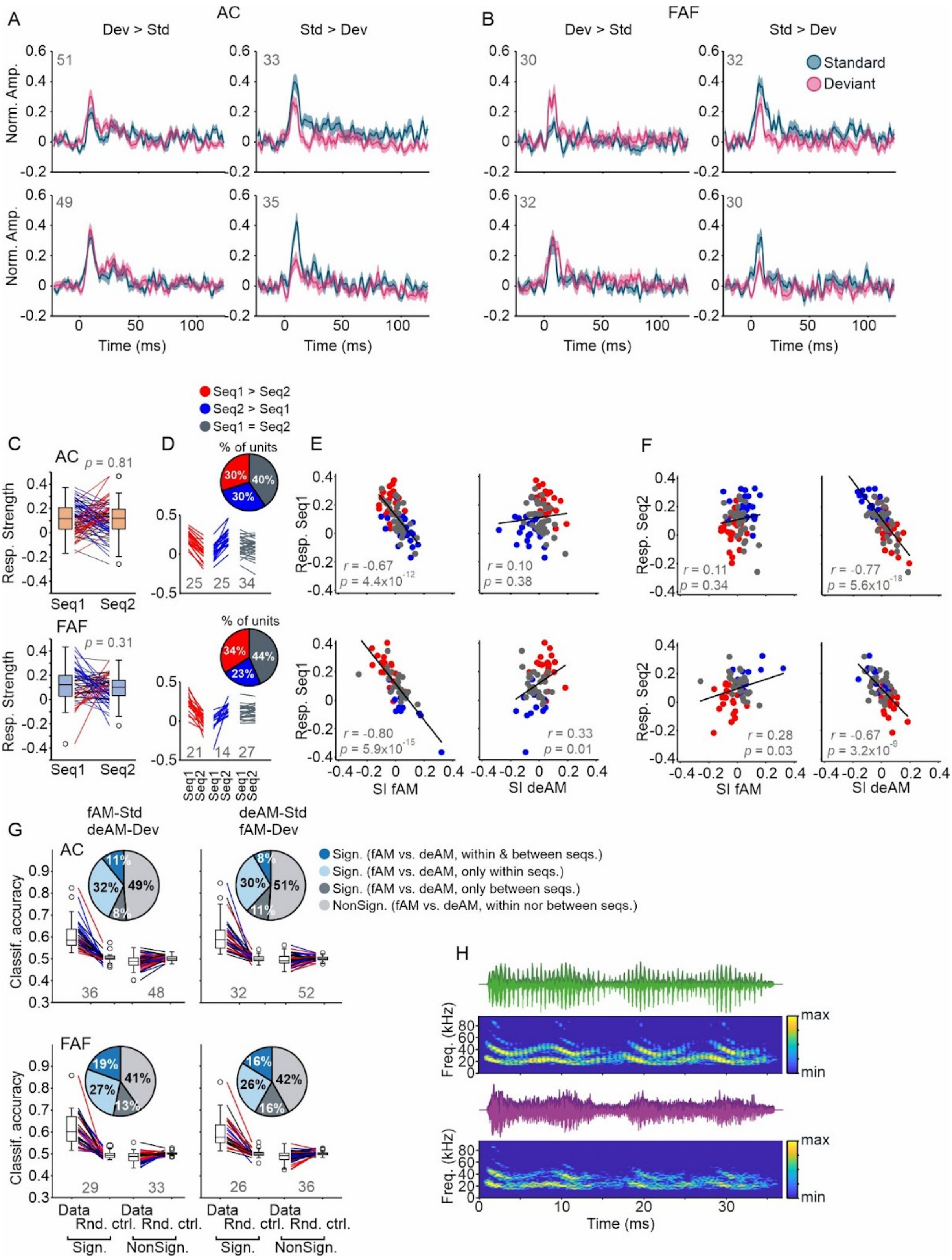


Figure S1. Neural activity with a natural AM call and its demodulated version. A-B) Mean ± SEM (standard error of the mean) of probability density functions (Gaussian kernel of width 1 ms, at steps of 0.5 ms) during the deviant (pink) responses to the sounds (fAM, top row; deAM, bottom row) and the standard (dark blue, using the same number of sound presentations as for the deviant N = 40 trials)

for the AC (A) and FAF (B). C) Response strength (mean of activity response in a sequence, considering 40 standards) for each sequence (Seq1: fAM as Standard; Seq2: deAM as Standard). There were no significant differences between the response in the two sequences (Wilcoxon signed-rank test) when all the units were considered. We further classified units into three groups, in red (significant response difference between Seq1 and Seq2, Wilcoxon rank-sum test, $p<0.05$, with response in Seq1 > Seq2), in blue (same as for the red group, but with response in Seq2 > Seq1), and gray (no significant difference between Seq1 and Seq2). Each line represents a unit, with the total number of units shown below. The plots are shown for the AC (top) and FAF (bottom). Insets show the percentage of units in each group. D) Line plots showing the same as in C, but for all groups separated. E) Scatter plots of the response strength in Seq1 and the SI of both stimuli fAM (left) and deAM (right) for AC (top) and FAF (bottom). Linear regression fit (gray line) for all units, with the Pearson's correlation coefficient and its corresponding *p* value also shown. E) Same as in D for Seq2. G) Box plots of the prediction accuracy calculated using binary SVM classifiers, same conventions as for Figure 4 of the main part of the manuscript. H) Oscillograms (in darker color, the envelope is shown; secant method, temporal resolution: 0.05 ms) and spectrograms (512 points, hamming window, 0.8 ms frame width and 0.05 ms frame shift).

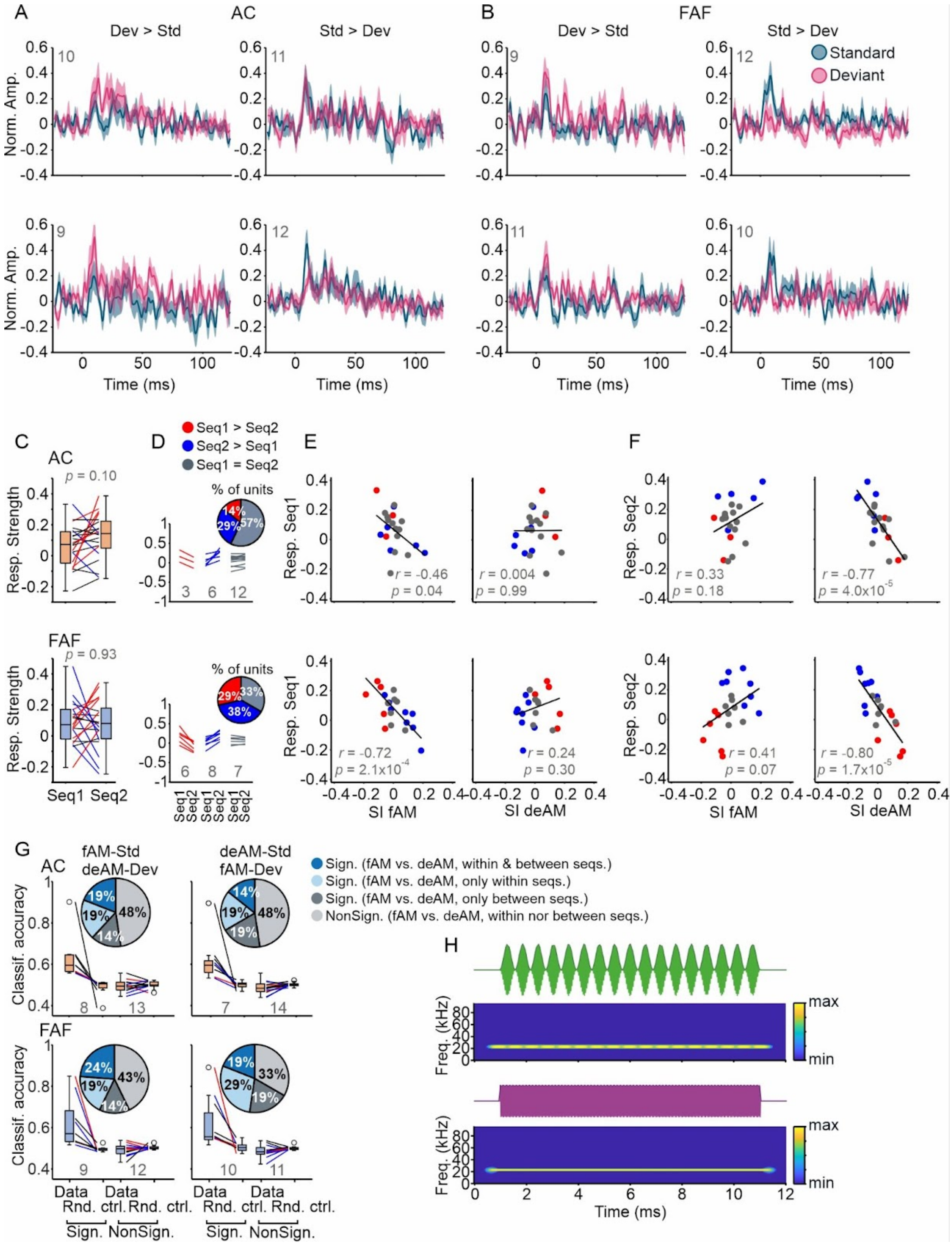


Figure S2. Neural activity with an AM and non-modulated pure tone (23 kHz). A-B) Mean ± SEM (standard error of the mean) of probability density functions (Gaussian kernel of width 1 ms, at steps of 0.5 ms) during the deviant (pink) responses to the sounds (fAM, top row; deAM, bottom row) and the standard (dark blue, using the same number of sound presentations as for the deviant N = 40 trials) for the AC (A) and FAF (B). C) Response strength (mean of activity response in a sequence, considering 40 standards) for each sequence (Seq1: fAM as Standard; Seq2: deAM as Standard).

There were no significant differences between the response in the two sequences (Wilcoxon signed-rank test) when all the units were considered. We further classified units into three groups, in red (significant response difference between Seq1 and Seq2, Wilcoxon rank-sum test, $p<0.05$, with response in Seq1 > Seq2), in blue (same as for the red group, but with response in Seq2 > Seq1), and gray (no significant difference between Seq1 and Seq2). Each line represents a unit, with the total number of units shown below. The plots are shown for the AC (top) and FAF (bottom). Insets show the percentage of units in each group. D) Line plots showing the same as in C, but for all groups separated. E) Scatter plots of the response strength in Seq1 and the SI of both stimuli fAM (left) and deAM (right) for AC (top) and FAF (bottom). Linear regression fit (gray line) for all units, with the Pearson's correlation coefficient and its corresponding *p* value also shown. E) Same as in D for Seq2. G) Box plots of the prediction accuracy calculated using binary SVM classifiers, same conventions as for Figure 4 of the main part of the manuscript. H) Oscillograms (in darker color, the envelope is shown; secant method, temporal resolution: 0.05 ms) and spectrograms (512 points, hamming window, 0.8 ms frame width and 0.05 ms frame shift).

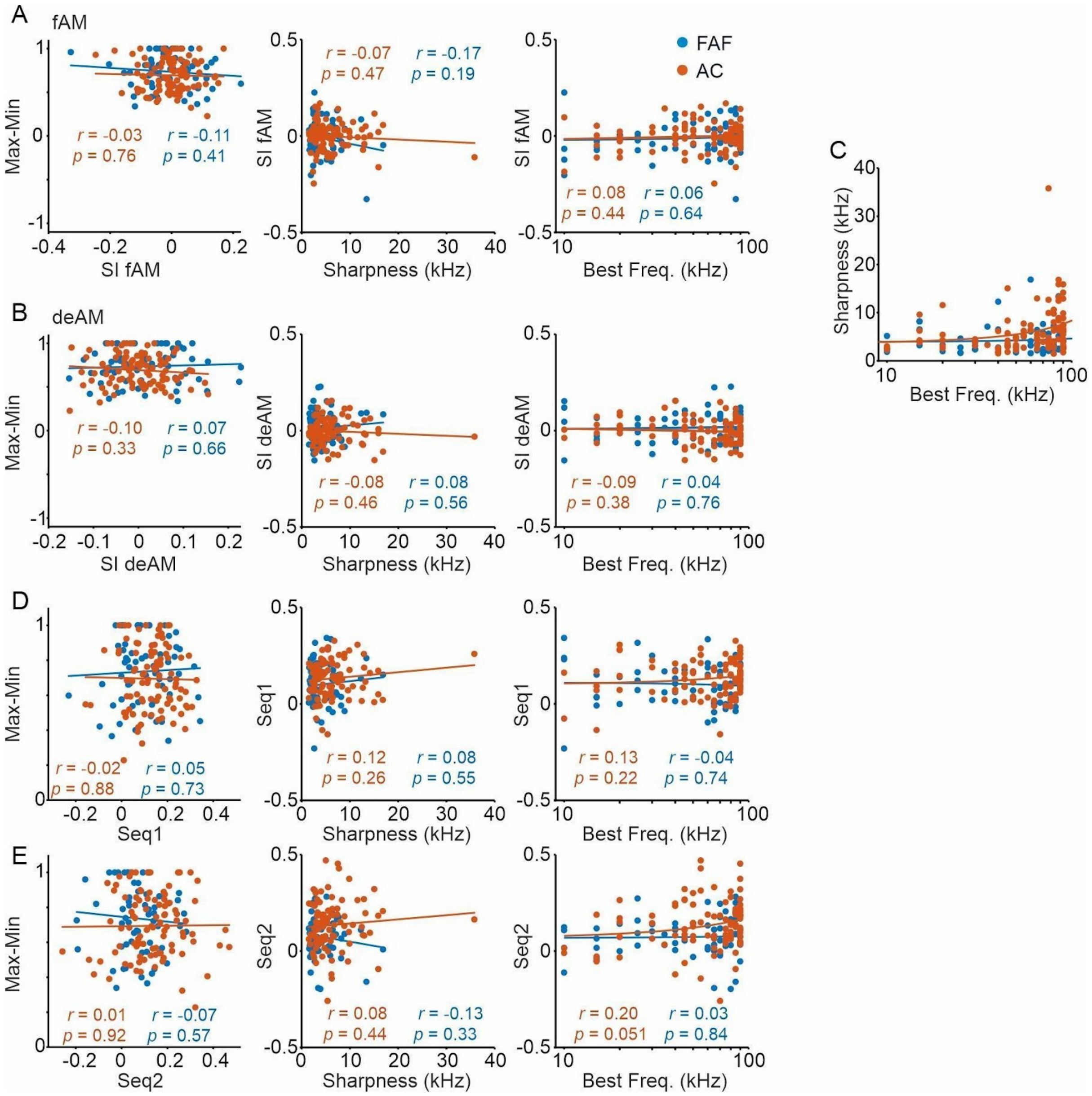


Figure S3. Frequency tuning properties. A) Left; SI of the stimulus fAM plotted against the difference between the maximum and minimum of the normalized frequency tuning curves (spike counts of the 110 ms since stimulus onset, pure tones played: from 10 to 90 kHz in steps of 5 kHz, at 70 dB SPL). Middle: sharpness (distance between the frequencies where a line at 85% of the maximum crosses the frequency tuning curve before and after the best frequency) plotted against the SI of fAM. Right: best frequency (frequency with the largest number of spikes). B) Same as in A, for the stimulus deAM. C) Best frequency plotted against the sharpness, with logarithmic x axis. Exponential fit for FAF: f(x)=3.886*exp(0.002*x), and AC: f(x)=3.613*exp(0.008*x). D-E) Same as A-B but considering the response strength during Sequence 1 (Seq1; fAM as Standard) and for Sequence 2 (Seq2; deAM as Standard). In blue, Frontal Auditory Field (FAF) and in orange, Auditory Cortex (AC).

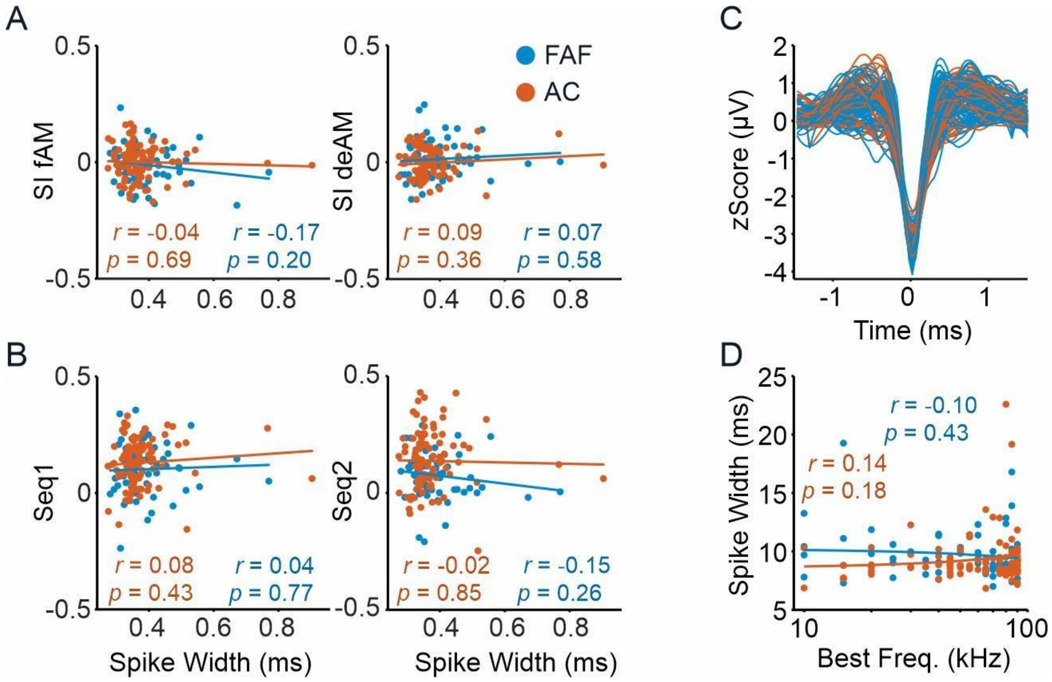


Figure S4. Spike shape. A) Scatter plot of the spike width (measured as the width at 0.75 of absolute of the Hilbert transform, normalized to its maximum value) plotted against the SI of both stimuli, fAM (left) and deAM (right) and for the Frontal Auditory Field (FAF, in blue) and Auditory Cortex (AC, in orange). Plotted is also the linear regression fit, the Pearson's correlation coefficient and its corresponding *p* value. B) As in A, with spike width plotted against the response strength for Sequence 1 (Seq1, fAM as Standard, deAM as Deviant, left) and for Sequence 2 (Seq2, deAM as Standard, fAM as Deviant, right). C) Normalized spike shapes of the units color-coded for the recording area (blue, FAF; orange, AC). D) Best frequency plotted against the spike width (logarithmic x axis).

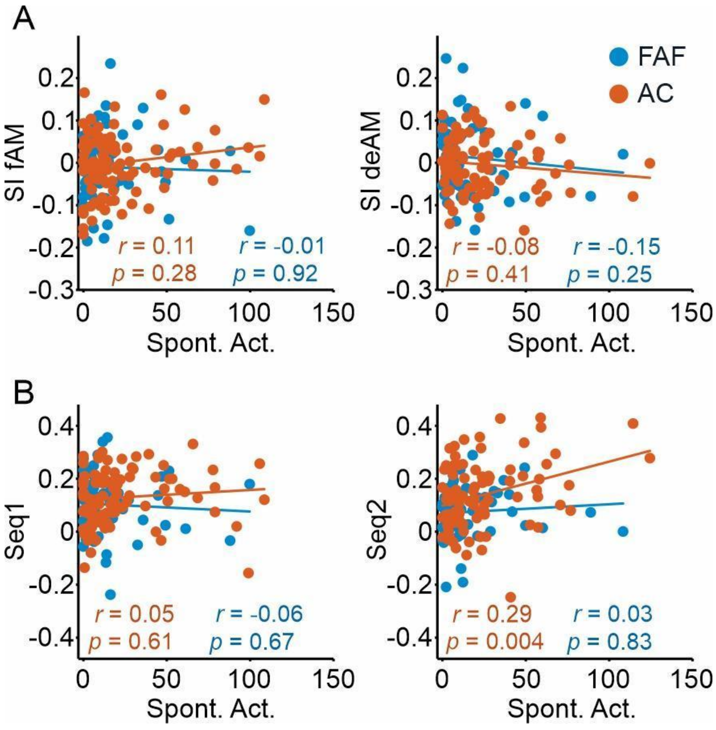


Figure S5. Spontaneous activity is only correlated with the responsivity of deAM in the Auditory Cortex. A) Scatter plot of the spontaneous activity (measured as the average spiking rate in 5 2-s windows before each stimulation sequence) plotted against the response strength for Sequence 1 (Seq1, fAM as Standard, left) and for Sequence 2 (Seq2, deAM as Standard, right), and for the Frontal Auditory Field (FAF, in blue) and Auditory Cortex (AC, in orange). Plotted is also the linear regression fit, the Pearson's correlation coefficient and its corresponding *p* value. B) As in A, showing the Spontaneous Activity plotted against the SSA Index (SI) of both stimuli, fAM (left) and deAM (right).

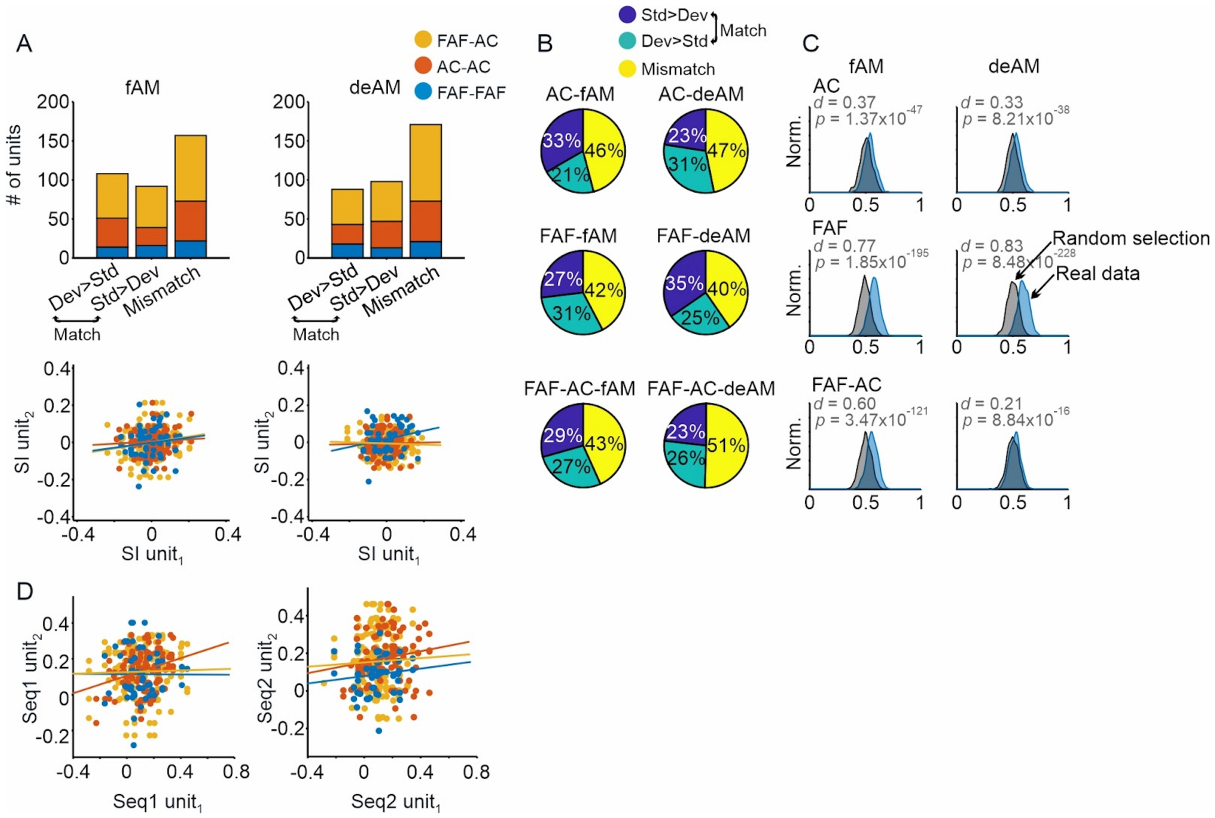


Figure S6. Simultaneously-recorded pairs. A) Histograms of pairs of simultaneously-recorded units per stimulus and group combination (top) and scatter plots of the SIs for each unit of the pair (bottom). Linear regression fits are also shown (non-significant Pearson's correlation coefficient). For fAM, AC-AC (Pearson's *r* correlation coefficient: 0.07, *p*=0.45), FAF-FAF (Pearson's *r* correlation coefficient: 0.12, *p*=0.40) and FAF-AC (Pearson's *r* correlation coefficient: 0.18, *p*=0.01), and for the deAM stimulus, AC-AC pairs (Pearson's *r* correlation coefficient: 0.02, *p*=0.83), FAF-FAF pairs (Pearson's *r* correlation coefficient: 0.23, *p*=0.10) and FAF-AC pairs (Pearson's r correlation coefficient: -0.04, *p*=0.58). B) Proportion of matching (both units "Dev>Std" or both "Std>Dev", see legend) and mismatching pairs. C) Probability density functions (Gaussian kernel of width 0.01, at steps of 0.01) of the probability of finding a matching pair (i.e. finding two units belonging to the same group) taking random units (black) or actual pairs from the real data (blue), considering 100 pairs in 1000 repetitions. Shown is also the Cliff's delta (*d*) and the *p* value of a Wilcoxon rank sum test between the two distributions. D) Scatter plots of the response strength (Seq1 for sequence with fAM as standard; Seq2 for the sequence with deAM as standard) for each unit of the pair. For Seq1, AC-AC (Pearson's *r* correlation coefficient: 0.28, *p*=0.003), FAF-FAF (Pearson's *r* correlation coefficient: -0.004, *p*=0.98) and FAF-AC (Pearson's *r* correlation coefficient: 0.03, *p*=0.73), and for the Seq2, AC-AC pairs (Pearson's *r* correlation coefficient: 0.15, *p*=0.11), FAF-FAF pairs (Pearson's *r* correlation coefficient: 0.12, *p*=0.40) and FAF-AC pairs (Pearson's *r* correlation coefficient: 0.04, *p*=0.54).

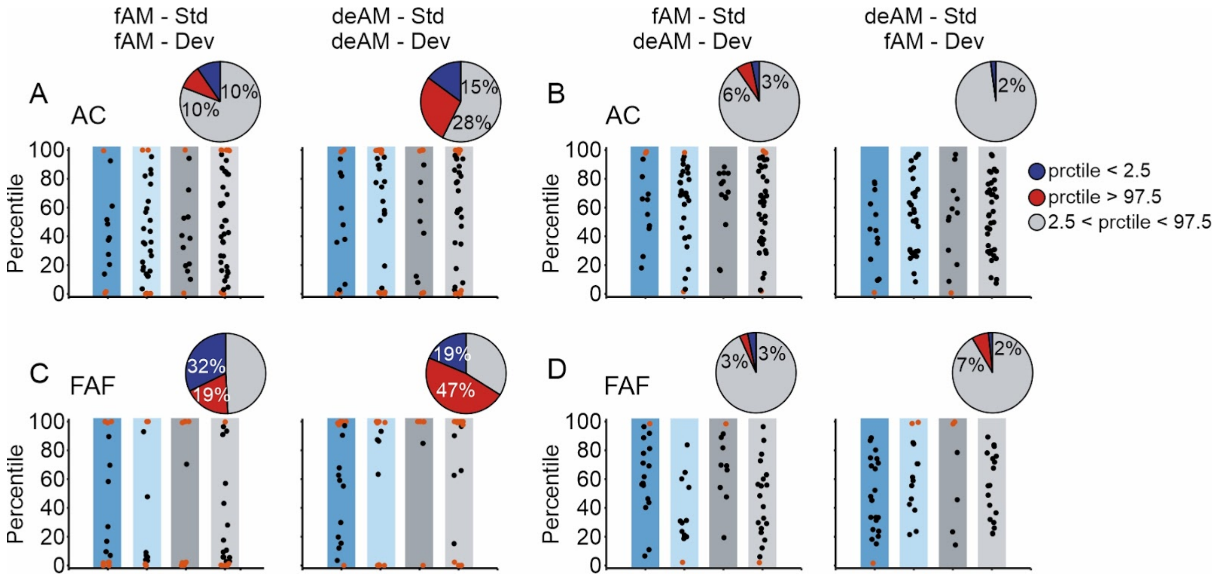


Figure S7. Percentiles of a bootstrapping test. Firing rate calculated for 40 random standard presentations (repeated 1000 times) during the first 25 ms after sound onset in order to create a distribution. The firing rate of the deviant (40 presentations) is compared to the distribution and the respective percentile is plotted. Orange dots if the percentile (prctile) is lower than 2.5 or higher than 97.5, otherwise black dots. Shown for the AC (A & B) and FAF (C & D), and for comparisons of the same sound in different sequences (A & C; as for the SI calculation) and for fAM and deAM in the same sequence (B & D). Units shown grouped according to the classification in Figure 4 of the main part of the manuscript. Pie charts show the percentage of significant (red and blue colors) and non-significant units in each panel (see legend).
